# Single-shot water-fat imaging with multi-echo blip-rewound EPI

**DOI:** 10.1101/2024.05.13.593911

**Authors:** Wenchuan Wu

## Abstract

**Purpose:** To develop a method for single-shot water-fat imaging.

**Methods:** Single-shot blip-rewound EPI (rEPI) is a fast imaging technique that can acquire multiple echo MRI images with a snapshot. Dixon’s method separates water and fat signals using signals acquired with different echo times. Combining multi-echo rEPI with properly chosen echo times and Dixon-based post-processing generates water and fat images with a single-shot scan.

**Results:** Phantom and in vivo experiments were performed to evaluate the proposed method. In phantom scans, water and fat signals are robustly separated using the proposed method, and the obtained water signal is much cleaner than that obtained by conventional EPI acquisition with fat suppression. In vivo experiments show that fat suppression can be obtained using the proposed method without fat saturation pulses, reducing SAR and shortening scan time. In addition, fat signal can also be reconstructed based on the proposed method.

**Conclusion:** A single-shot water-fat imaging technique has been successfully developed, a significant advancement that offers speed and robustness in separating water and fat signals for human MRI scans.

## Introduction

Conventional MRI scans detect signals from protons, which may come from different molecules, including water and fat. Due to its very short T1 relaxation time, fat signals typically appear very bright in the image, which can obscure underlying pathologies. Particularly, with EPI-based acquisition, fat signals can be shifted along the phase encoding direction and overlap with water signals, resulting in chemical shift artefacts. Therefore, fat suppression has been an important research area in MRI for several decades.

Conventional fat suppression methods include: a) chemically selective fat suppression^1^, b) spatial-spectral pulse based excitation (i.e., water only excitation) ^2,3^, c) short inversion time inversion recovery imaging ^4,5^, and d) chemical-shift based water-fat separation ^6^. Chemically selective fat suppression (a) and water only excitation (b) methods are sensitivity to B0 field inhomogeneity. Chemically selective fat suppression methods (a) are also sensitive to transmit field (B1+) inhomogeneity. Short inversion time inversion recovery imaging (c) would introduce mixed contrast in the water signal and has a low SNR efficiency. In comparison, only chemical-shift based water-fat separation methods are robust against B0 and B1+ inhomogeneity and does not introduce unwanted imaging contrast, which are ideal for robust fat suppression. In addition, chemical-shift based water-fat separation can also provide quantification of fat signals, which can be potentially used to quantify disease severity like hepatic steatosis and other diseases due to abnormal fat deposition.

Chemical shift-based water-fat separation requires the acquisition of multiple images with different echo times, which are typically achieved by multiple shots. However, this approach would result in long scan times and higher sensitivity to motion. Motion-induced inconsistencies between shots may introduce unwanted phase and magnitude variations that may degrade the accuracy of the downstream water-fat separation. Hence, a rapid imaging method with a single-shot acquisition can improve the robustness of water fat imaging against motion.

In this work, a single-shot based water-fat imaging method is developed. This method builds on the recently developed single-shot blip-rewound EPI^7^ to acquire multi-echo images in a single readout, and calculates separate water and fat images based on Dixon’s technique^6^. Preliminary data from phantom and in vivo experiments demonstrate capability of the proposed method for rapid and robust water-fat imaging.

## Methods

### Blip-rewound EPI

Blip-rewound EPI (rEPI) is a new EPI-based acquisition technique that is able to acquire multiple echoes in a single readout^7^ (Fig. 1). Compared to conventional single-shot EPI, rEPI trajectory has a zig-zag pattern in the ky-t space, which samples adjacent lines (each line corresponds to one echo) in an interleaved order (Fig. 1a). In k space, rEPI traverses along the phase encoding direction with a periodic rewinding pattern, enabling interleaved acquisition of multiple echoes (Fig. 1b). The implementation of rEPI trajectory is straightforward. Fig. 1c illustrates an example of spin-echo rEPI. Compared to conventional spin-echo single-shot EPI, the spin-echo single-shot rEPI sequence inserts rewinder blips between standard EPI blips. Previously, rEPI has been demonstrated for dynamic B0 mapping and distortion correction with gradient echo preparation^7^. Here, spin-echo based rEPI will be used for single-shot water fat imaging. The acquired rEPI data can be split into multiple k-space, each corresponding to an under-sampled conventional EPI k-space with a different echo time (Fig. 1d). The multiple echo images can be separately reconstructed using conventional parallel imaging reconstruction or jointly reconstructed to leverage shared information between echoes^7^.

**Figure 1.**
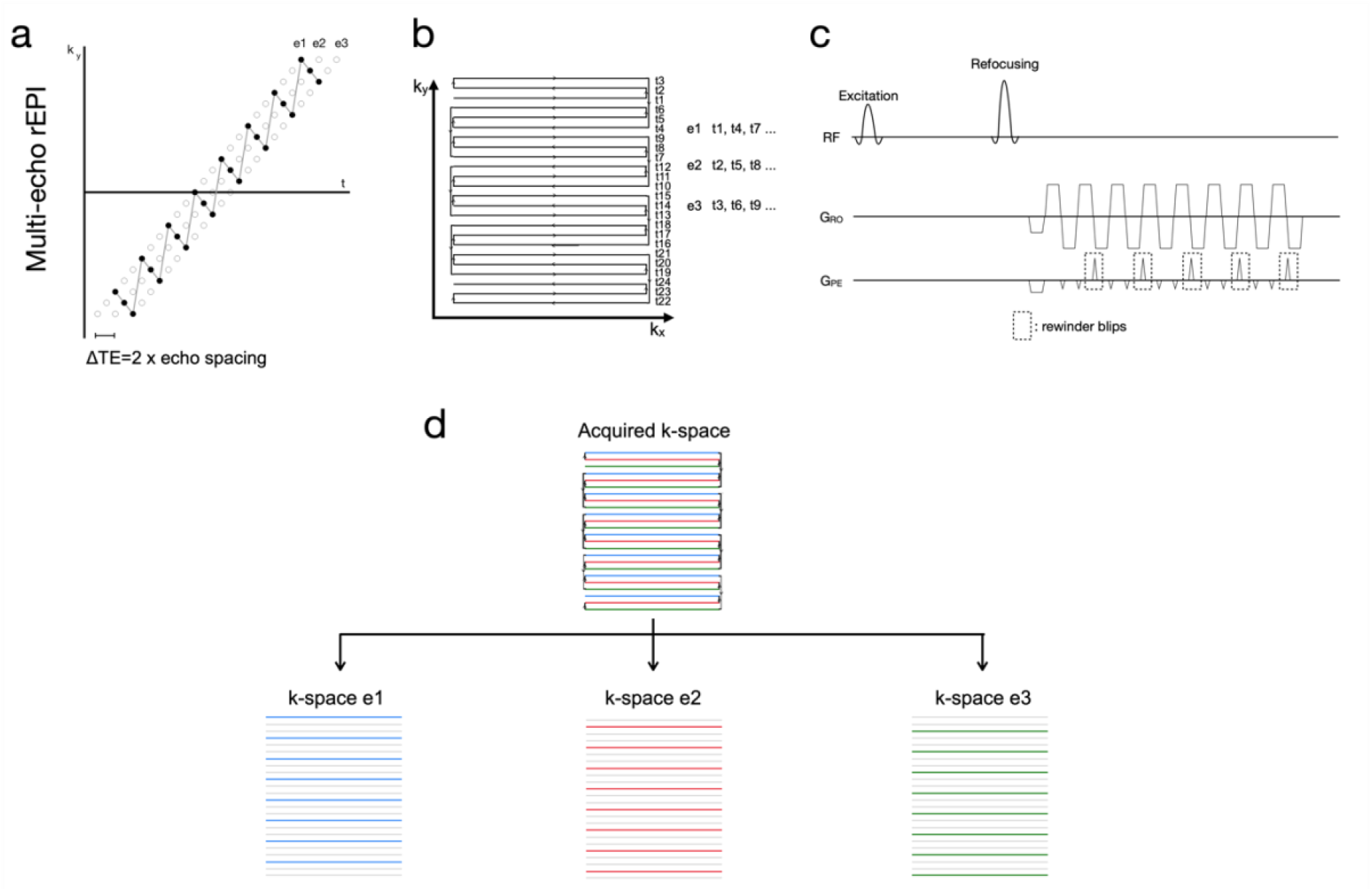
rEPI trajectory in ky-t space (a), kx-ky space (b) and implementation diagram (c). (d) The rEPI data can be split into multiple k-space, each corresponding to a conventional single-shot EPI k space with a different echo time.

### Dixon water-fat separation

The basic concept of Dixon’s method is illustrated in figure 2 using two echoes as an example. Due to the different precessional frequencies of water and fat, signals can be acquired with water and fat signal in-phase and out of phase. This is achieved by manipulating the echo time (TE) such that the accumulated phase difference between water and fat is π × *even _number* for in-phase scans and π × *odd_number* for out-of-phase scans. Then water-only and fat-only signals can be obtained using simple calculation. For more robust water-fat separation using Dixon type methods, three-point Dixon method was developed^8^, which acquires three images with specific TE values that produce a phase shift of +π, 0 and −π between the water and fat signals. The three-point Dixon addressed the problem of B0 inhomogeneity, making it a more robust option. Iterative decomposition of water and fat with echo asymmetry and least square estimation (IDEAL) technique was also developed for water fat separation ^9^, which allows for three or more echoes at arbitrary echo times (i.e., no need to generate exactly +π, 0 and −π phase shift between water and fat).

**Figure 2.**
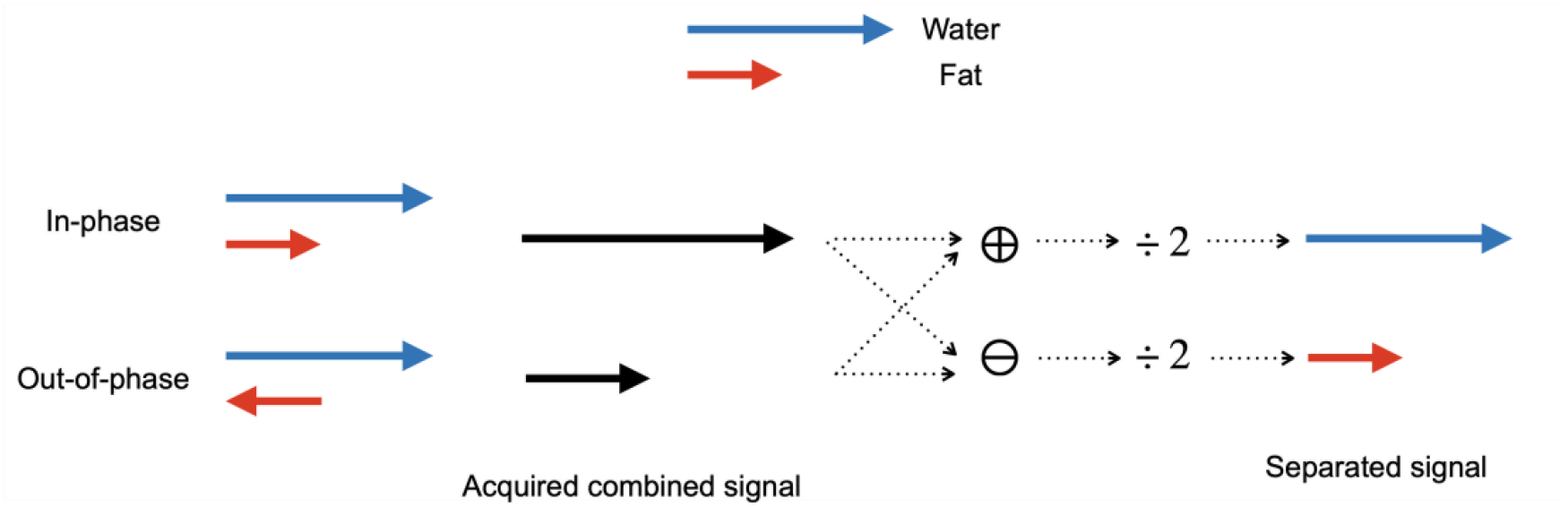
Basic concept of Dixon’s method. In two-echo Dixon method, two images with different TEs are acquired, and the water signal and fat signal are in-phase in one image (i.e., signal vectors of water and fat are aligned with each other, thus, total signal = water signal + fat signal) and out-of-phase in the other image (i.e., signal vectors of water and fat are pointing opposite directions, thus, total signal = water signal - fat signal). By adding or subtracting the in-phase image and out-of-phase image, separate water and fat images can be obtained.

The proposed method was implemented. Phantom data and in vivo data from one subject were acquired to evaluate the performance of the proposed method on a 3T whole body MRI scanner (Siemens Prisma, Erlangen, Germany) using a 32-channel receive coil.

### Phantom experiment

Data was acquired using a spherical fat phantom and a water tube phantom (positioned in the same FOV). Three datasets were acquired: conventional single-shot EPI without fat saturation, conventional single-shot EPI with fat saturation (vendor provided ‘Fat Sat.’), and single-shot 3-echo rEPI without fat saturation. Scan parameters for conventional single-shot EPI: FOV 220mmx220mmx100mm, resolution 2.2mmx2.2mmx2mm, TR=7s, echo spacing 0.65ms, no in-plane acceleration applied, phase encoding direction is left-right. Scan parameters for rEPI are the same as conventional single-shot EPI, but with a shorter TR=6.4s. The echo time difference between two consecutive echoes in rEPI is 1.3ms.

The center frequency was manually set to the frequency of the water during the scan.

### In vivo experiment

One subject was scanned to evaluate the performance of rEPI based water-fat separation. Three datasets were acquired: conventional single-shot EPI without fat saturation, conventional single-shot EPI with fat saturation (vendor provided ‘Fat Sat.’), and single-shot 3-echo rEPI without fat saturation. The scan parameters are the same as in the phantom experiment. Written informed consent in accordance with local ethics was obtained before the scan.

## Results

A spherical oil phantom and a water tube were positioned in the same FOV in the phantom experiment. In Figure. 3, both water and fat signals are visible using conventional single-shot EPI without fat suppression. Using conventional single-shot EPI with vendor-provided Fat Sat., the fat signal is partially suppressed and the residual fat signal is still strong, which is likely due to the severe B0 inhomogeneity. Using rEPI based water-fat separation, the water-only signal is much cleaner than that using conventional EPI with fat suppression. Also, with rEPI acquisition and Dixon-based water-fat separation, we can obtain an image of fat.

**Figure 3.**
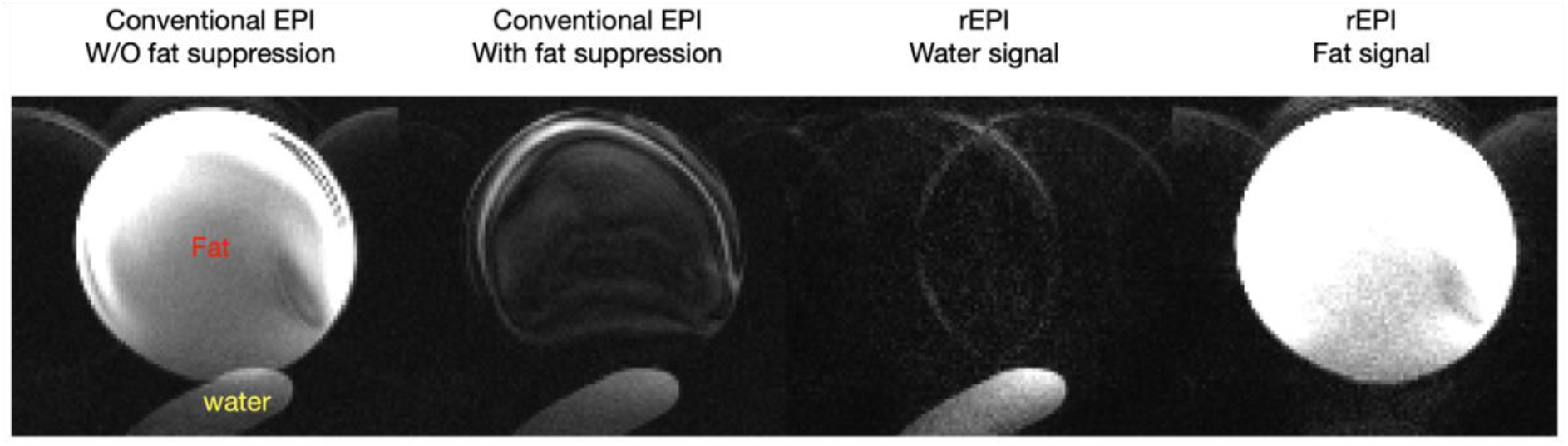
Phantom experiment comparing conventional EPI with fat suppression and rEPI based fat-water separation. A spherical oil phantom and a water tube were imaged in the same FOV. From left to right: conventional EPI without fat suppression, where both fat and water signal are visible; conventional EPI with fat suppression, where fat signal is just partially suppressed due to limitations of the conventional fat suppression technique (i.e., sensitivity to B1 and B0 inhomogeneity); water image from rEPI based water-fat imaging; fat image from rEPI based water-fat imaging.

Figure 4 compares conventional single-shot EPI with fat suppression with rEPI-based water fat separation from a subject scan. With rEPI-based water-fat separation, water-only images can be acquired without using a fat saturation pulse, reducing SAR requirement and shortening scan time. By integrating rEPI and Dixon’s method, the fat signal can also be obtained using a single-shot scan.

**Figure 4.**
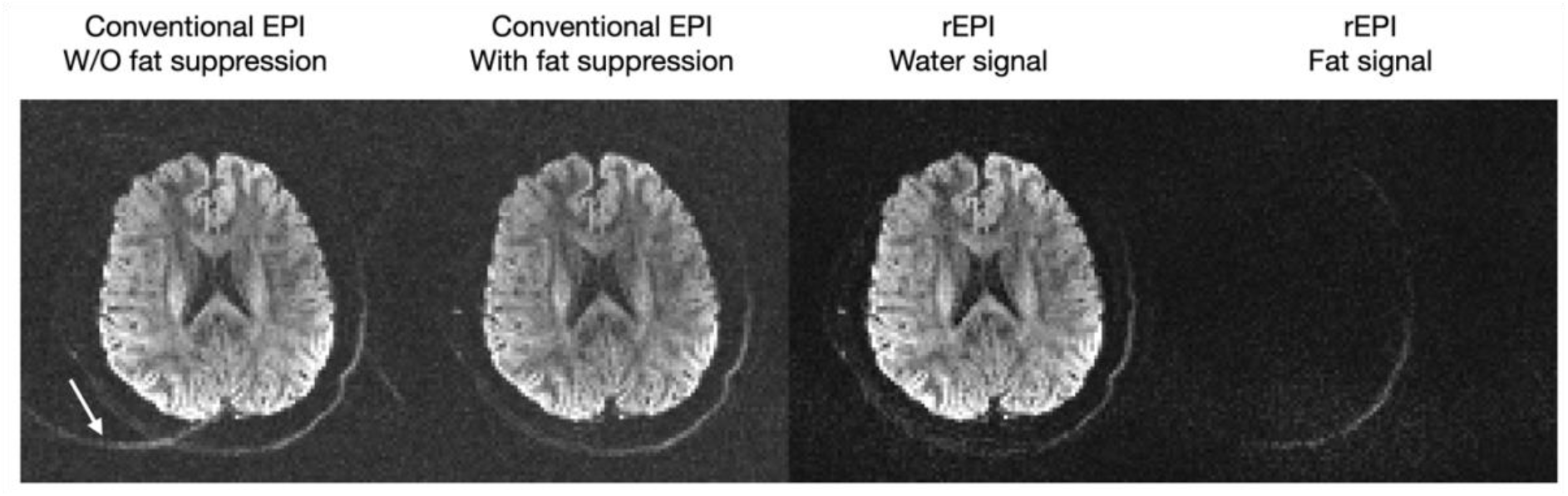
In vivo experiment comparing conventional EPI with fat suppression and rEPI based fat-water separation. From left to right: conventional EPI without fat suppression, where both fat and water signal are visible. Due to the low bandwidth along the phase encoding direction (left-right), the fat signal is shifted relative to the water signal (white arrow); conventional EPI with fat suppression, where fat signal is suppressed; water image from rEPI based water-fat imaging; fat image from rEPI based water-fat imaging.

## Discussion and conclusion

In this work, a method for single-shot water fat imaging is developed. This method integrates multi-echo rEPI trajectory with the Dixon technique to achieve single-shot water-fat imaging. Preliminary results from phantom and in vivo subject demonstrate the feasibility of the proposed method.

Here, rEPI data were acquired without in-plane acceleration, which was used to test the worst-case scenario, where the fat signal can be shifted significantly along the phase encoding direction, introducing overlapping with the water signal in the image. The proposed method can robustly separate fat signal and water signal, as shown in the phantom and in vivo results. For this protocol, individual echo signal (under-sampling factor R=3) was reconstructed using conventional parallel imaging GRAPPA ^10^. However, for in-plane under-sampled data, joint multi-echo reconstruction is needed, as demonstrated previously^7^.

The phantom experiment demonstrates a scenario where conventional fat saturation breaks down due to extreme B0 inhomogeneity. Nevertheless, using the proposed method, we can still get very clean water and fat images, demonstrating the robustness of this method against B0 inhomogeneity.

Here, the key concept of integrating rEPI and Dixon method for single-shot water fat imaging is tested as a proof of principle. Future work will investigate the proposed method’s capability in more challenging tasks, such as imaging moving organs where single-shot measurement would be more beneficial for motion-robust water fat imaging.

## Reference

1. Haase A, Frahm J, Hanicke W, Matthaei D. 1H NMR chemical shift selective (CHESS) imaging. Phys Med Biol. 1985;30(4):341. doi:10.1088/0031-9155/30/4/008

2. Schick F. Simultaneous highly selective MR water and fat imaging using a simple new type of spectral-spatial excitation. Magnetic Resonance in Medicine. 1998;40(2):194–202. doi:10.1002/mrm.1910400205

3. Meyer CH, Pauly JM, Macovskiand A, Nishimura DG. Simultaneous spatial and spectral selective excitation. Magnetic Resonance in Medicine. 1990;15(2):287–304. doi:10.1002/mrm.1910150211

4. Bydder GM, Pennock JM, Steiner RE, Khenia S, Payne JA, Young IR. The short TI inversion recovery sequence—An approach to MR imaging of the abdomen. Magnetic Resonance Imaging. 1985;3(3):251–254. doi:10.1016/0730-725X(85)90354-6

5. Bydder GM, Steiner RE, Blumgart LH, Khenia S, Young IR. MR Imaging of the Liver Using Short TI Inversion Recovery Sequences. Journal of Computer Assisted Tomography. 1985;9(6):1084.

6. Dixon WT. Simple proton spectroscopic imaging. Radiology. 1984;153(1):189–194. doi:10.1148/radiology.153.1.6089263

7. Dynamic field mapping and distortion correction using single-shot blip-rewound EPI and joint multi-echo reconstruction - Wu - 2024 - Magnetic Resonance in Medicine - Wiley Online Library. https://onlinelibrary.wiley.com/doi/full/10.1002/mrm.30038. Accessed May 13, 2024.

8. Glover GH, Schneider E. Three-point dixon technique for true water/fat decomposition with B0 inhomogeneity correction. Magnetic Resonance in Medicine. 1991;18(2):371–383. doi:10.1002/mrm.1910180211

9. Reeder SB, Pineda AR, Wen Z, et al. Iterative decomposition of water and fat with echo asymmetry and least-squares estimation (IDEAL): Application with fast spin-echo imaging. Magnetic Resonance in Medicine. 2005;54(3):636–644. doi:10.1002/mrm.20624

10. Griswold MA, Jakob PM, Heidemann RM, et al. Generalized autocalibrating partially parallel acquisitions (GRAPPA). Magnetic Resonance in Medicine. 2002;47(6):1202–1210. doi:10.1002/mrm.10171

